# AI-assisted In-silico Trial for the Optimization of Osmotherapy following Ischaemic Stroke

**DOI:** 10.1101/2024.07.20.604439

**Authors:** Xi Chen, Lei Lu, Tamás I. Józsa, David A. Clifton, Stephen J. Payne

## Abstract

Over the past few decades, osmotherapy has commonly been employed to reduce intracranial pressure in post-stroke oedema. However, evaluating the effectiveness of osmotherapy has been challenging due to the difficulties in clinical intracranial pressure measurement. As a result, there are no established guidelines regarding the selection of administration protocol parameters. Considering that the infusion of osmotic agents can also give rise to various side effects, the effectiveness of osmotherapy has remained a subject of debate. In previous studies, we proposed the first mathematical model for the investigation of osmotherapy and validated the model with clinical intracranial pressure data. The physiological parameters vary among patients and such variations can result in the failure of osmotherapy. Here, we propose an AI-assisted in-silico trial for further investigation of the optimisation of administration protocols. The proposed deep neural network predicts intracranial pressure evolution over osmotherapy episodes. The effects of the parameters and the choice of dose of osmotic agents are investigated using the model. In addition, clinical stratifications of patients are related to a brain model for the first time for the optimisation of treatment of different patient groups. This provides an alternative approach to tackle clinical challenges with in-silico trials supported by both mathematical/physical laws and patient-specific biomedical information.

## I. Introduction

HE proper functioning of the human brain relies on a continuous and sufficient blood flow that sustains cerebral metabolism. In the context of an ischaemic stroke, blood vessel blockage can cause a reduction in cerebral blood flow (CBF) and blood-brain barrier (BBB) breakdown, resulting in a rise in BBB permeability [1, 2]. Reperfusion therapy aims at restoring CBF to the affe**c**ted area and can thus elevate blood pressure and lead to excessive fluid leakage into the interstitial space (see Fig. 1). Consequently, this fluid accumulation in the interstitial space can result in brain oedema, increasing intracranial pressure (ICP), and excessive brain tissue stress [3]. Elevated ICP leads to symptoms including headache, nausea, vomiting, and loss of consciousness. Typically, a value of ICP exceeding 25 mmHg is deemed life-threatening and necessitates immediate medical intervention [4]. Over the past few decades, osmotherapy has been widely employed to restore ICP to the normal range of 5 to 15 mmHg [4, 5].

**Figure 1:**
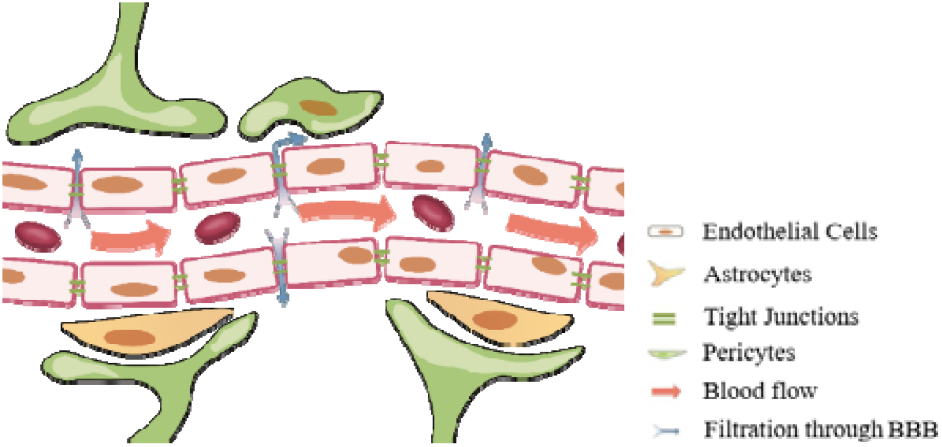
A schematic of fluid filtration from the lumen to the interstitial space through the blood brain barrier (BBB) of a single capillary vessel.

Currently, four types of osmotic agents (mannitol, hypertonic saline, sorbitol, and glycerol) are available for treatment and hypertonic saline has gained increasing popularity [6, 7]. Hypertonic saline creates an osmotic pressure gradient that drives excessive fluid from the brain into the intravascular space, thereby reducing the volume of interstitial fluid and relieving oedema. Despite its various advantages, the use of osmotic agents can result in a variety of issues [7, 8]. One concern is that the elevation in Cl^-^ level in hypertonic saline can lead to hyperchloremic acidosis [9]. Additionally, a high concentration of NaCl (exceeding 160 mmol/L) is reported to worsen outcomes and further damage the blood-brain barrier [9, 10]. Hence, it is crucial to carefully select the administration protocol. In clinical practice, despite the fact there have been previous comparisons of osmotherapy outcomes [11-13], no optimised administration protocol has yet been established for hypertonic saline infusion. The variations in the proposed protocols in previous studies have been hypothesised to be a result of heterogeneity in the patients [2].

To tackle this issue, the current study aims to expand upon simulation tools developed in the In Silico Clinical Trials for the Treatment of Acute Ischaemic Stroke (INSIST) project [14, 15]. The objective is to incorporate AI-assisted in-silico osmotherapy trial into the existing Finite Element model (FEM) and explore treatment protocols that are optimal in reducing ICP. The deep learning model can incorporate statistics and therefore establish a connection between patient characteristics and low information-noise ratio clinical data using an FEM model based on mathematical and mechanical theories. Once a reliable model has been established, it can enhance the precision of resource-intensive clinical studies, consequently reducing the enormous associated costs. Drawing inspiration from recent studies on biofluid transport in the brain [16-20], recent AI applications in biomedical studies [21-23], and the application of AI in the field of engineering [24, 25], the first AI-assisted in-silico trial for the investigation of osmotherapy is thus proposed here.

In this model, we will present the prediction of the ICP evolution using DNN model during osmotherapy episodes on a quasi-population level. The study consists of the following parts: (1) the FEM model and the training and testing of the neural network will be presented; (2) the effects of physiological parameters on the main features of the osmotherapy will be investigated; (3) the effectiveness of the osmotherapy will be evaluated using two criteria and the effects of different physiological parameters will be presented; (4) proper doses of osmotic agents will be proposed for different levels of BBB damage and age

## II. Methods

The model consists of two components, including the FEM model and the artificial neural network. In the previously developed FEM model [26], the brain is assumed to consist of four compartments, i.e., tissue, arteriole, capillary, and venule network and can thus be considered a multi-network fluid dynamic system. The modelling pipeline consists of three 3D models for the simulation of blood perfusion, oedema, and osmotherapy, respectively. After in-silico trials, the DNN model is trained using the ICP data generated from the FEM model.

### A. Perfusion Model

#### 1) Governing Equations

In the perfusion component, the brain model consists of arteriole, venule, and capillary blood compartments, with each compartment a porous medium [18, 26, 27]. The pial surface is subdivided into perfusion regions and the perfusion in stroke scenarios can be obtained by occluding perfusion territories (Fig. 2a) and compared with healthy perfusion. The governing equations of the three-compartment perfusion model are taken directly from our previous studies:

**Figure 2:**
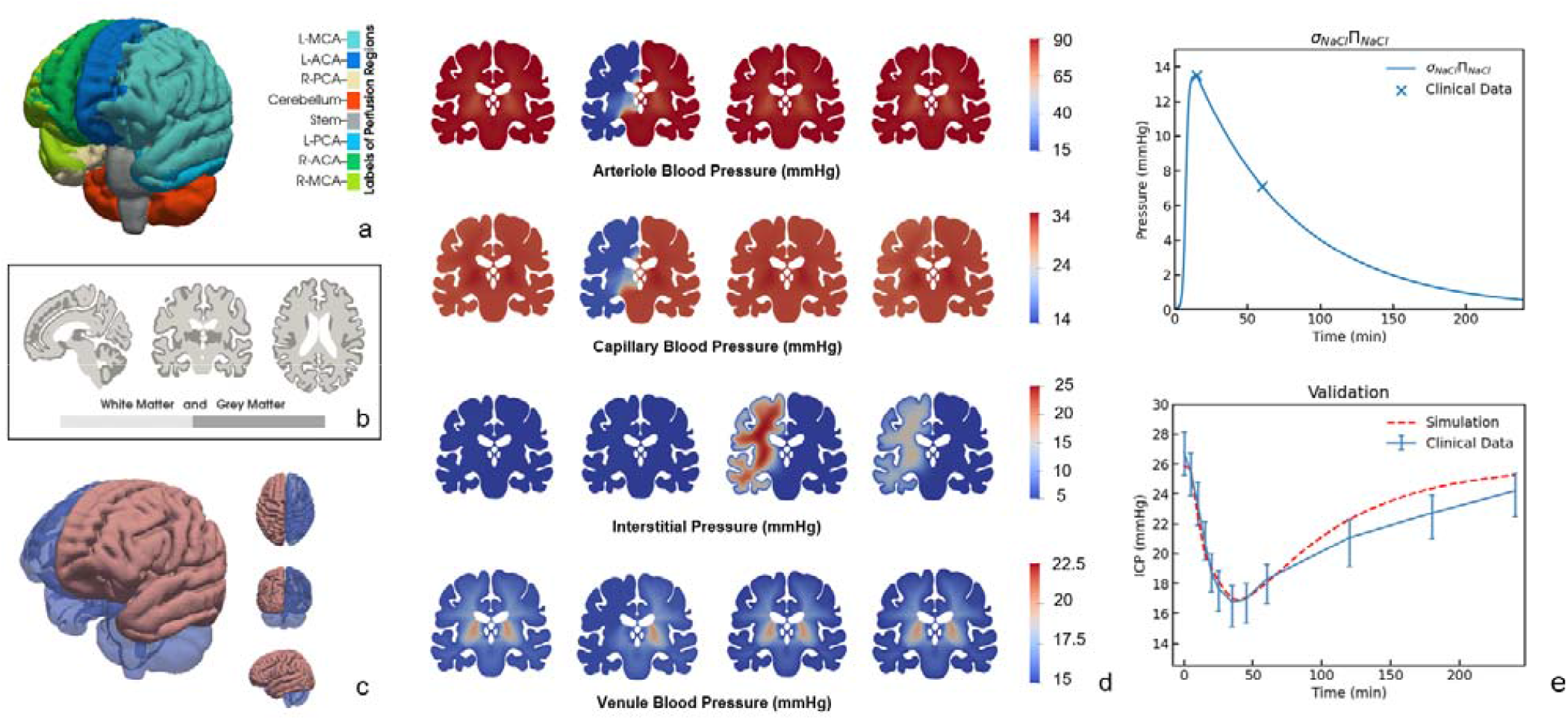
(a) Superficial perfusion territories corresponding to large arteries, where L-PCA, L-MCA, L-ACA and R-PCA, R-MCA, R-ACA represent groups. the posterior, middle and anterior cerebral arteries on the left and right hemisphere, respectively. The perfusion territories can be occluded to obtain the occluded blood perfusion. (b) white and grey matter in the brain model. (c) Left hemisphere oedema volume generated using the perfusion model. (d) blood pressure and interstitial pressure along the coronal plane, from left to right: healthy, stroke, oedema, and osmotherapy. (e) Comparison of the model with clinical osmotic pressure and ICP data, reproduced from Chen et al., 2022, without change.

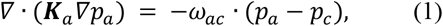

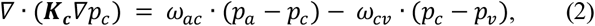

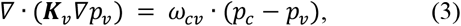

where, and are the Darcy pressures in the arteriole, capillary, and venule compartments respectively. is the permeability tensor of compartment, where the capillary network tensor field can be replaced with the scalar field. ω_*ij*_ represents the fluid transfer coefficient between compartments *i*and *j*. The transfer coefficients are different in grey matter and white matter (Fig. 2b) and are therefore further defined by 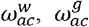 and 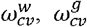.

#### 2) Boundary Conditions

Blood feeds the brain through penetrating arterioles and flows back into the veins through penetrating venules at the pial surface. Therefore, only arteriole and venule compartments have non-zero blood flow at the pial surface. In the model, a 90mmHg arteriole blood pressure and a 15mmHg venule pressure are given on the pial surfaces. According to a vascular territory atlas [28], each artery feeds one sub-territory at the pial surface, as shown in Fig. 2a. Note that blood flow through the perfusion territory of an occluded vessel is set to zero whereas pressure on the pial surface is assumed to remain constant in other regions. The boundary conditions of the perfusion model are given in Table 1.

**TABLE 1.**
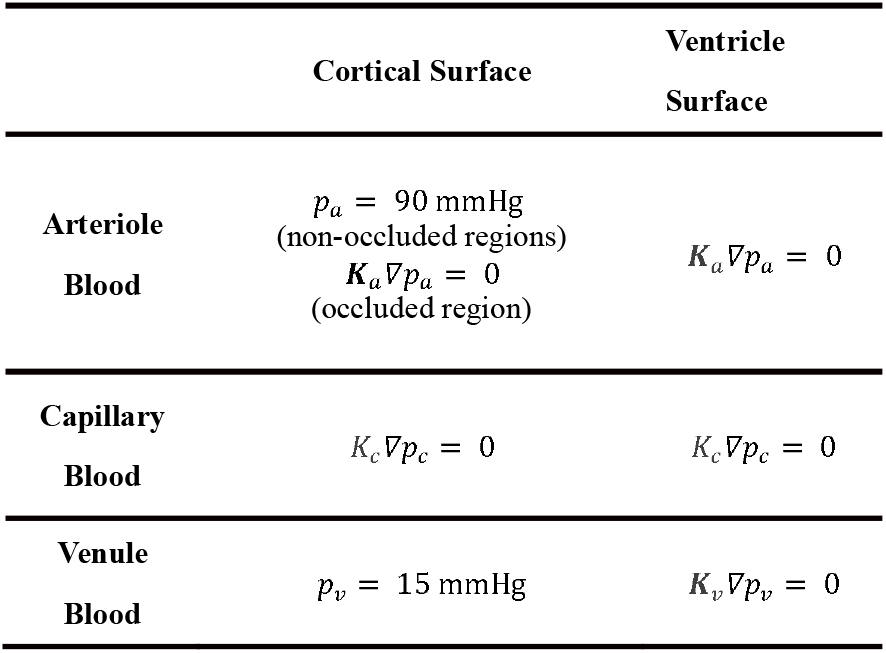
Boundary conditions for the perfusion model.

### B. Oedema and Osmotherapy Model

#### 1) Governing Equations

The flow exchange between the capillary network and the interstitial space is negligible in a healthy brain. However, in the context of a damaged brain, an extra compartment is necessary for interstitial fluid in both the oedema and osmotherapy models to simulate the ICP change. Furthermore, to model ICP evolution in oedema and osmotherapy, time-dependent terms are introduced to the governing equations:

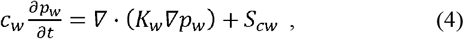

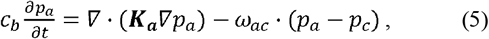

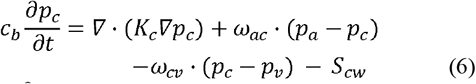

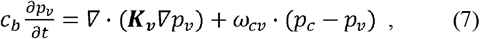

where *C*_*w*_ and *C*_*b*_ are the storage factors of the interstitial fluid and blood in the brain tissue. The storage factors in the blood compartments are assumed to be the same. The parameters *p*_*w*_ and *k*_*w*_ are the pressure and hydraulic conductivity of interstitial fluid in the interstitial space. The *S*_*cw*_ is the fluid transport from the vasculature into the interstitial space. Note that in the equations above, we neglect any terms related to displacement of the tissue; this is done as the time scale for pressure variations is much longer than that for the displacement variations [29]. Hence, the temporal derivatives of the displacement terms can be neglected at the time scale which we are considering here. *S*_*cw*_ is derived in Appendix A and summarised in Table 2.

**TABLE 2.** Fluid transfer through the capillary wall in the oedema and osmotherapy model.

|  | Oedema | Healthy Tissue |
| --- | --- | --- |
| **Oedema Model** | $S_{cw} = 2n_b \frac{L_p}{R_c} [(p_c - p_w) - \sigma \Pi_c]$ | 0 |
| **Osmotherapy Model** | $S_{cw} = 2n_b \frac{L_p}{R_c} [(p_c - p_w) - \sigma \Pi_c - \sigma_{NaCl} \Pi_{NaCl}]$ | 0 |

*L*_*p*_ is the hydraulic permeability of the capillary wall, *Π*_c_ is the osmotic pressure for the plasma components in the blood before osmotherapy, σ is the reflection coefficient of the original plasma composition, *n*_*b*_ is the volume fraction of blood vessels in a unit volume of brain tissue, and *R*_c_ is the mean vessel radius. In osmotherapy, the osmotic pressure rise induced by the hypertonic saline is described by an additional term σ_*NaCl*_ Π_*NaCl*_ where σ_*NaCl*_ is the reflection coefficient of hypertonic saline and the Π_*NaCl*_ osmotic pressure of the saline. Similarly, due to the large difference in the BBB permeability between damaged and healthy tissue, the fluid transfer through the capillary wall can be neglected in healthy tissue. The corresponding simulation results are shown in Fig. 2. Here, the expression [26] for the effective osmotic pressure is taken to be:

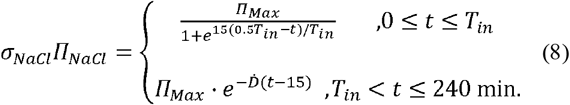

where the infusion time *T*_*in*_ is fixed to be 15 min, and the decay rate 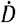 is set to be 0.01424min-1 to match the clinical osmotic pressure data (Fig. 2e) [26, 30]. Π_*Max*_ maximum osmotic pressure during an episode.

### 2) Boundary Conditions

In the oedema model, reperfusion therapy is performed, therefore it is assumed that there is no occlusion at the pial surface. Normal ICP is around 5-15 mmHg, and the baseline value of the interstitial fluid pressure is taken here to be 5 mmHg. The boundary conditions are as given in Table 3.

**TABLE 3.**
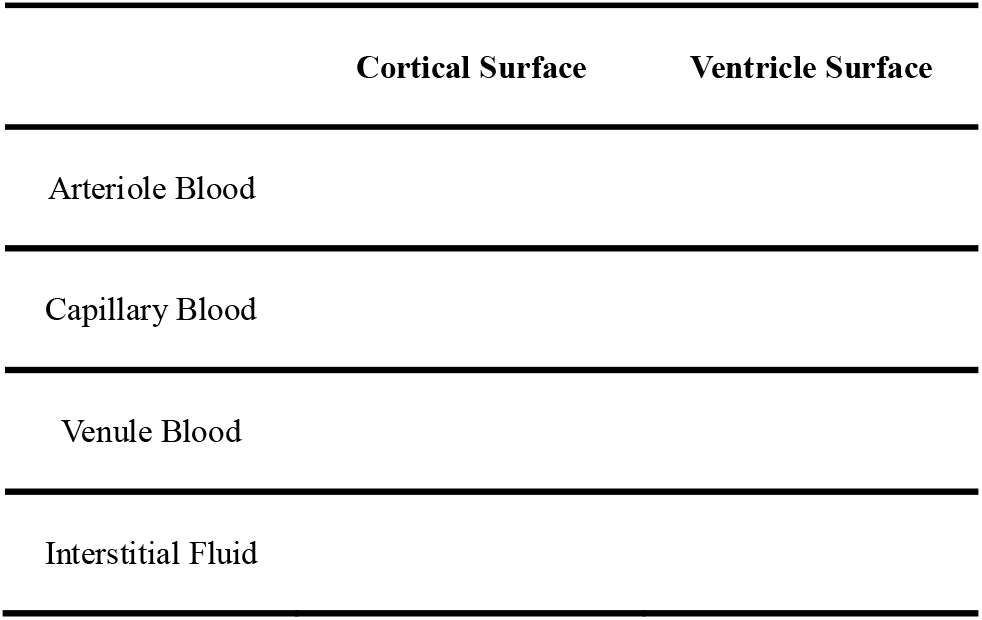
Boundary conditions for the oedema and osmotherapy model.

|  | Cortical Surface | Ventricle Surface |
| --- | --- | --- |
| Arteriole Blood |  |  |
| Capillary Blood |  |  |
| Venule Blood |  |  |
| Interstitial Fluid |  |  |

#### C. Design of Deep Neural Network Model

To facilitate the simulation process, we developed a deep neural network (DNN) model to explore the relationship between the parameters and the ICP change. The model has an input of five parameters, including the tissue storage factor C, tissue hydraulic conductivity, blood vessel permeability, arterial blood pressure ABP, and the maximum osmotic pressure. The parameters, and are varied between 0.5 to 2 times their baseline values, as these are the ranges that can be seen in previous studies [31-34], and the blood pressure ABP is varied between 80 mmHg to 100 mmHg [35]. Meanwhile, the maximum osmotic pressure is 1800 Pa in 10% hypertonic saline injection [26]. In the model, the maximum osmotic pressure is varied between 900-3600 Pa, which corresponds to the concentration (5%-20%) of hypertonic saline injection in literature [2, 36]. The output is a sequence of 49 data points for the ICP values at different time during an osmotherapy episode.

We tried different architectures for the DNN model design, including the models using dense layers, convolutional layers (Conv), long short-term memory (LSMT) layers, and residual neural network (ResNet). More advanced DNN models, such as transformer-based models [37], and diffusion models [38], have also been powerful tools for the regression prediction of biomedical data. In this study, however, we found that the ResNet neural network is sufficient for the prediction analysis of time series ICP data with a reasonably small loss. Using trial-and-error, we searched for an optimal model with the minimal error for the ICP value estimation. As shown in Fig. 3, the final searched DNN model has six convolutional layers and two ResNet blocks. In the first ResNet block, the convolutional layers have 48 kernels with the size of 3; and in the second ResNet block, the convolutional layers have 64 kernels with the size of 3. Then, a dense layer is used to produce the output for the ICP value estimation. The mean squared error between the ICP value and the estimation, summed over all time points, is used as the loss function. The model was trained for 100 epochs with a batch size of 16 and a learning rate of 0.001. Generally, the best model was decided based on the minimal loss during training.

**Figure 3:**
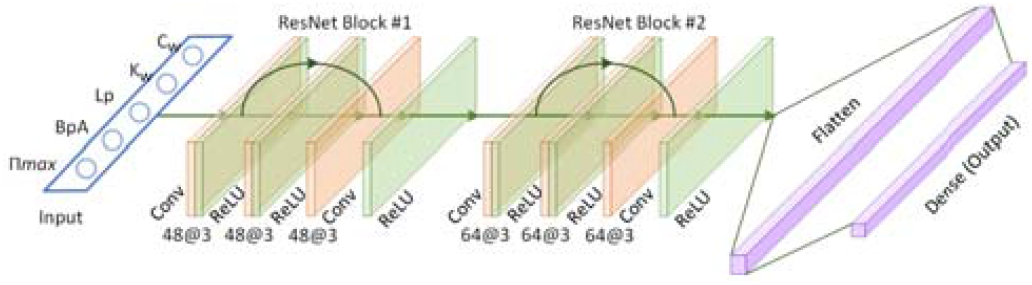
The architecture of DNN model for the ICP value prediction.

#### D. Model Coupling

The location of leaking vessels obtained from the perfusion model is then used within the oedema model. The perfusion () and perfusion change () are given as

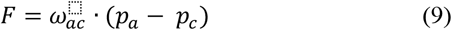

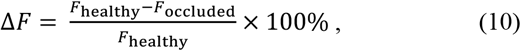

A threshold of 70% or greater blood perfusion reduction has been used in clinical imaging to determine brain damage regions [39, 40] and the threshold is thus utilised to obtain the geometry of oedema lesion in our model (Fig. 2c). Thereafter, the pressure field in the oedema is simulated until the ICP (defined to be the spatial maximum of) reaches equilibrium by using the stroke simulation results as the initial condition. Finally, the osmotherapy is assumed to be performed after the ICP reaches equilibrium and thus the pressures at the final step in the oedema model are used as the initial pressures in the osmotherapy model (Fig. 2d). A full space searching is performed for {, ABP,,, }, and 2000 in-silico trials were simulated. The data were then used to train and test the artificial neural network, using the process shown in Fig. 4. The model was validated with 8 patients ICP data who had 22 episodes from [30] (Fig. 2e) and the parameter sensitivity test is conducted by [26]. The baseline parameters are chosen to be the same as in the work of Chen et al. [26] (Table C1).

**Figure 4:**
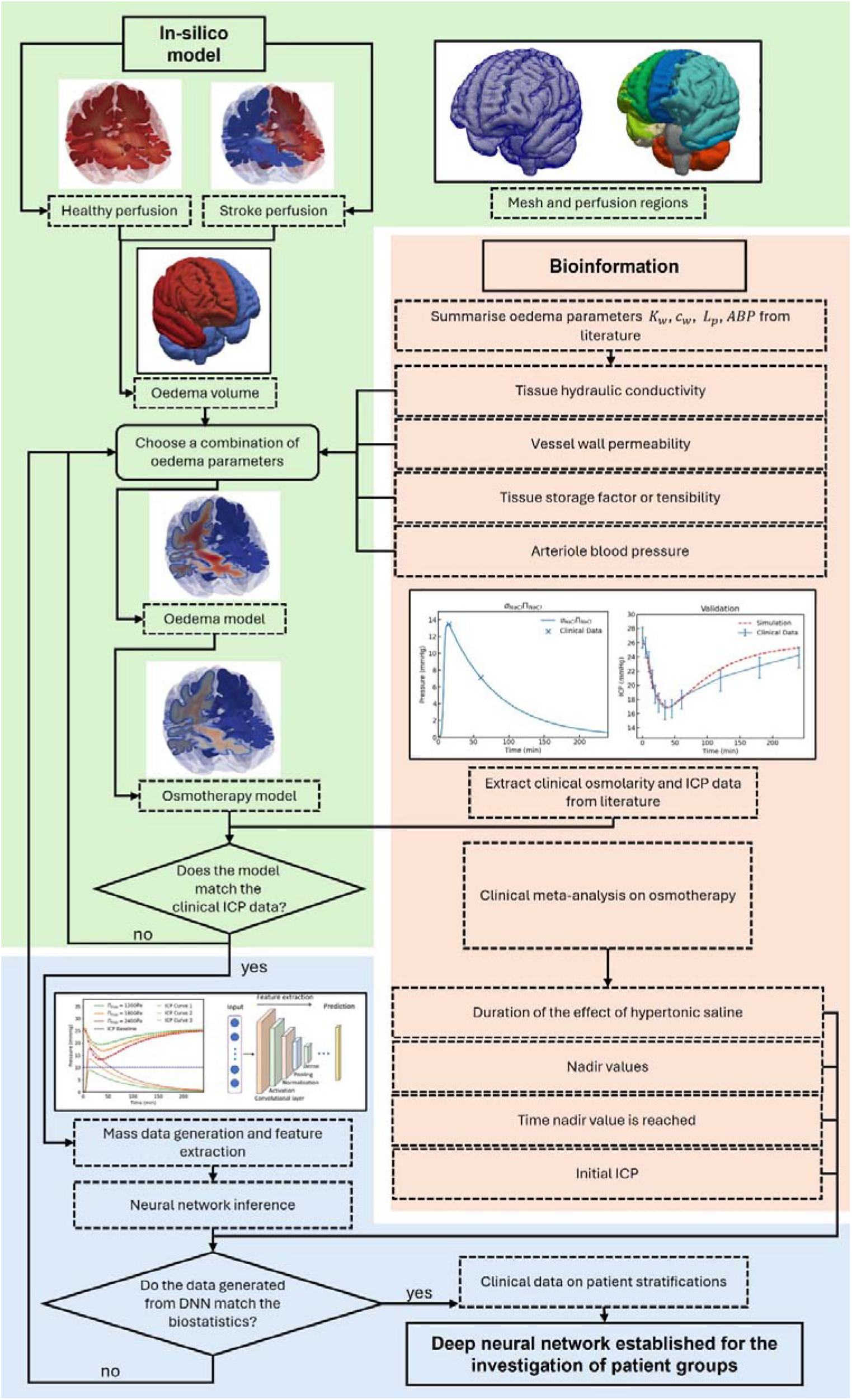
Flowchart demonstrating how the Finite Element models and bioinformation are coupled and used to generate data for deep neural network.

In the model, the governing equations are solved numerically using Python with a high-performance open-source Finite Element library, FEniCS [41]. The mesh used in this study has 1.4 million tetrahedral elements and the grid convergence study in the model has been performed in our previous study [19]. The DNN model is designed with Tensorflow 2.2, Python 3.7.4, and the Nvidia Quadro RTX 6000 is used for the model training. The inference of trained DNN is found to be faster than the conventional method and can thus be used for prediction of osmotherapy in large population.

## III Results

### A. DNN Model Training and ICP Value Estimation

We used a total of 2,000 in-silico samples for the DNN model development, out of which we randomly selected 1,600 samples for model training, and the remaining 400 samples were held out for model testing. To obtain a reliable model, we used 10-fold cross-validation to train the DNN model. We divided the whole dataset into training and testing sets, the training data were then randomly split into 10 folds. During each training, 9 of the 10 folds were used for model training and the rest fold was used for validation. For each iteration, the DNN model was trained for 100 epochs, and the model with the minimal estimation error was stored. The searched optimal model was then used to predict the ICP values in the hold-out testing dataset, which had the performance of 0.249 0.009 mmHg for the error of estimating ICP.

### B. Effect of Patient-specific Parameters on ICP Curve Evolution

We first investigate the effects of the physiological parameters on the features of ICP curves over an osmotherapy episode. The parameters vary with patients and are challenging to measure in clinical settings. However, previous studies have indicated a variation of the parameters in different patient groups. For example, the cerebrospinal fluid (CSF) clearance rate in smokers was reported to be lower than in non-smokers, indicating a lower tissue hydraulic conductivity [42]. Meanwhile, the blood vessel leakage was found to be larger in elderly patients, implying a larger value of [43]. In this study, we investigate the effects of different parameters by fixing one of the four parameter inputs and fixing the injection dose input while the rest of the parameters are set to be uniform random variables. The value of the,, or range from 0.5 to 2 times the baseline value and the ABP ranges from 80 mmHg to 100 mmHg. 2000 results are generated for each parameter using the DNN and the initial ICP, nadir ICP, nadir ICP time, and the duration of effect (defined as 10% ICP drop) are plotted as boxplots to show the difference on a simulation quasi-population level (Fig. 5).

**Figure 5:**
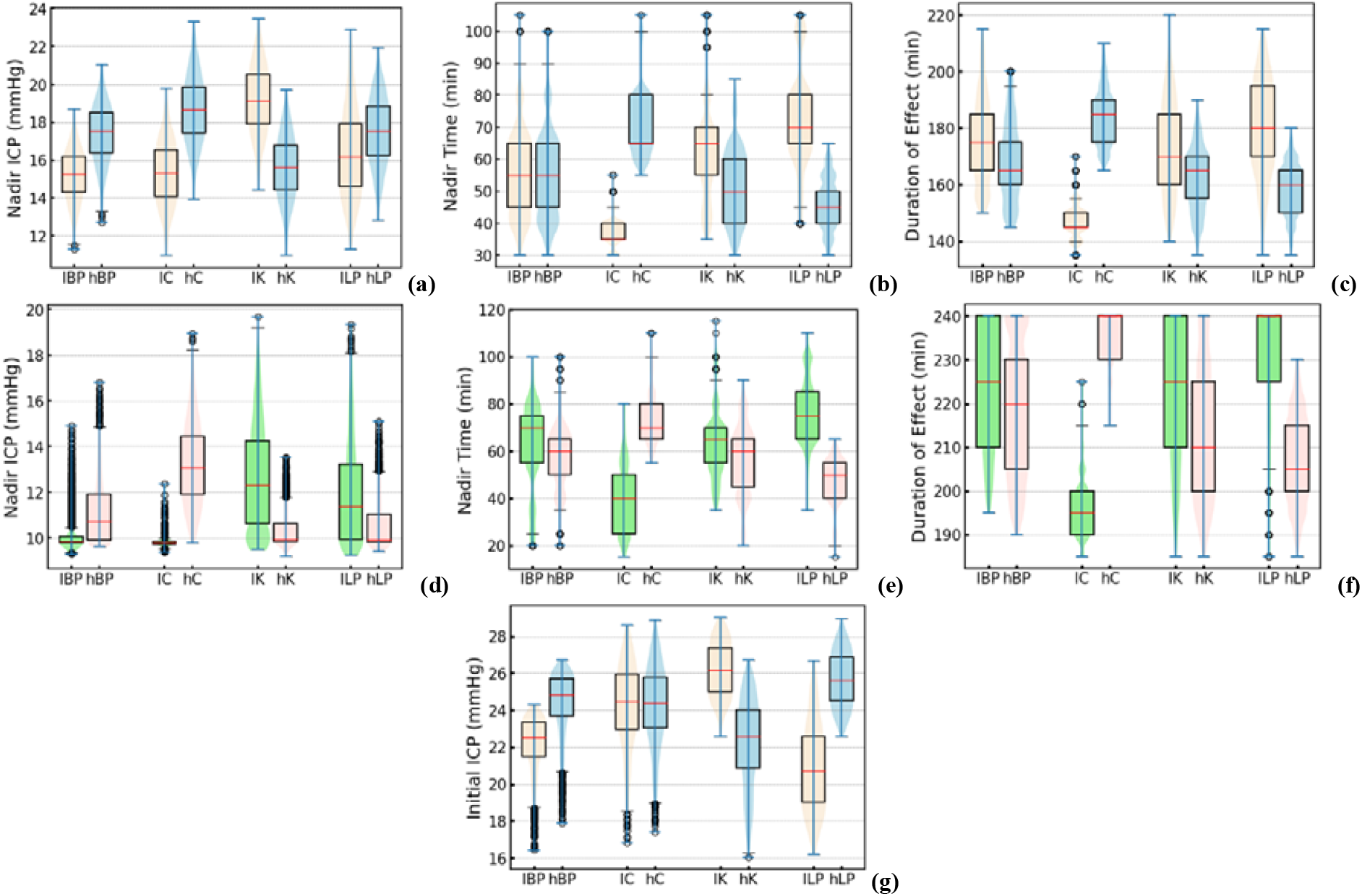
The features of the ICP curve as the flow parameters change. The first row shows the case where dose of injection leads to a maximum osmotic pressure of 1800 Pa, whereas the second row represents the case where maximum osmotic pressure is 3600 Pa. The third row shows the initial ICP. lBP, hBP denote 80 mmHg and 100 mmHg arteriole blood pressure, respectively. Meanwhile, the lC, hC, lK, hK, lLp, hLp represents 0.5 and 2 times the baseline values of the physiological parameters, with “l” for 0.5 and “h” for 2 times respectively.

The plots show that the and have the most significant effect on the initial ICP, with a mean of 20.8 to 25.6 mmHg and 22.6 to 26.1 mmHg, respectively (Fig. 5g). In contrast, the blood pressure value has only a limited effect on the initial ICP in the range around 22.8 to 24.7 mmHg. As to the nadir ICP values, it is noted that in both 1800 and 3600 Pa maximum osmotic pressure cases, the and presents a strong influence on the lowest ICP whilst the effects of the become less significant compared to its impact on initial pressure (Fig. 5a&d). Similarly, the ABP only shows a slight influence on the lowest pressure. Meanwhile, the time point nadir values are reached are most dependent on and, with a range from 20-115 mins. The effect of osmotherapy lasts from 135 to more than 240 mins.

### C. Evaluation of the Osmotherapy over an Episode Based on Two Criteria

Apart from the effects of the parameters on the ICP curves, the estimation of the effectiveness of osmotherapy is critical in designing a proper treatment plan for different patients. The evaluation of the effectiveness of osmotherapy has been limited by the difficulty in measuring ICP. In some studies, the evaluation is done via measuring the ICP nadir value or the ICP at certain time during an episode. For example, a 10% ICP drop 10 minutes after the end of infusion has been used to determine a successful therapy [30]. Alternatively, the pupillary diameter has also been used to estimate the drop in ICP [44]. Here, two criteria are utilised to estimate the effectiveness of the episode in the model. As the major goal of osmotherapy is to reduce ICP below 20 mmHg, the duration of time of the ICP below 20 mmHg in an episode is divided by the duration of the episode to obtain a time ratio and to indicate how effectively the ICP is reduced below this critical threshold. In addition, the overall pressure drop is divided by the overall rise in the osmotic pressure in the blood during an episode of 240 mins, as shown in Equation 11:

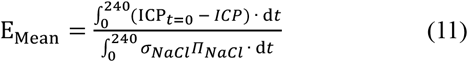

In Fig. 6, the maximum osmotic pressure here is chosen to be 1800 Pa. Virtual patient groups are generated, and 2000 episodes outcome are predicted in each group. To avoid cases with initial pressures below 20 mmHg, only those virtual patients with initial ICP over 25 mmHg are selected. The influences the both the efficiency and time ratio by around 15%, whereas the arteriole blood pressure shows a minor effect on efficiency but a significant impact on time ratio by 25%. Meanwhile, the storage factor and the time ratio by 17.2% and 16.1%, respectively.

**Figure 6:**
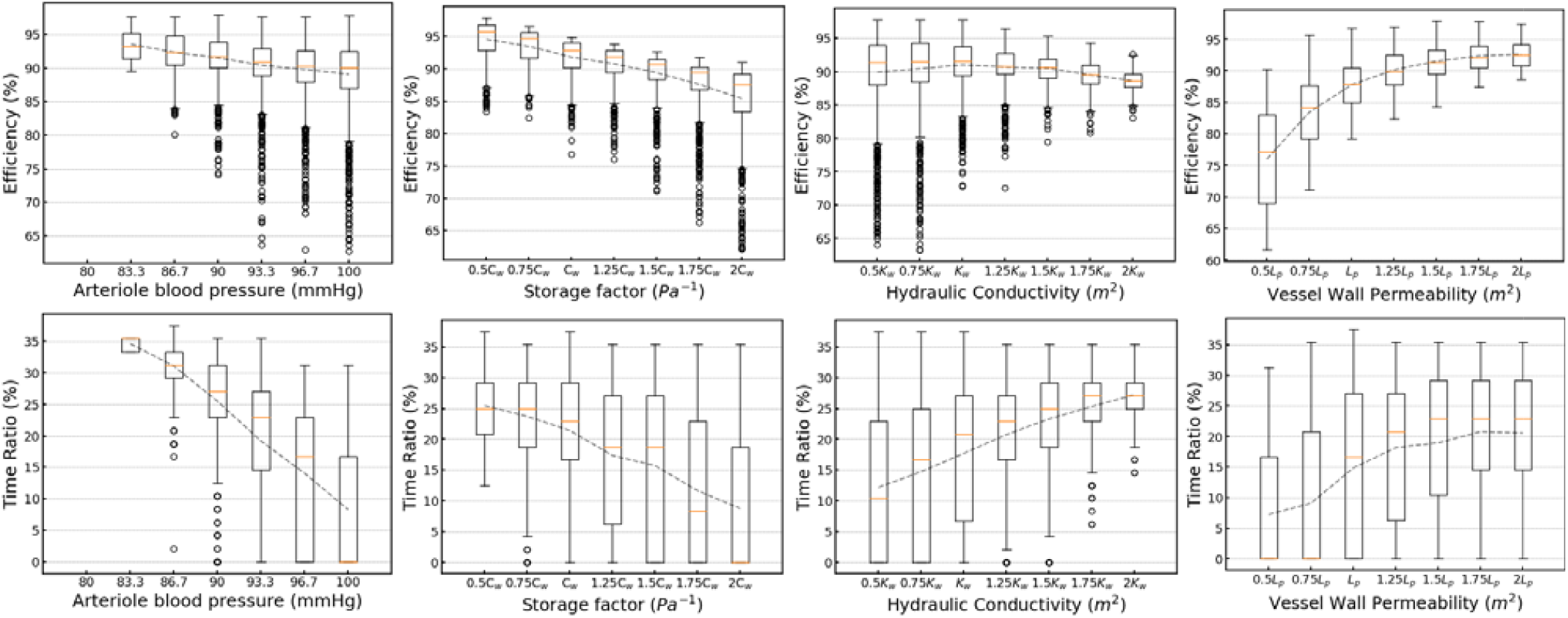
The treatment efficiency and the time ratio of ICP below 20 mmHg of an osmotherapy episode, where arteriole blood pressure (shows the blood pressure increment from baseline value of 80 mmHg, C is the storage factor, K is the tissue hydraulic conductivity and vessel wall permeability. The ABP’s baseline value is 80 mmHg with a uniform increment to the maximum of 100 mmHg. The boxplots show the mean and outliers in the quasi-population level trial and the dotted lines shows the average values of the different groups.

### D. The Optimization of Medication Doses for Various Brain Damage Levels

In clinical settings, the physiological parameters vary in different patient groups, especially the vessel wall permeability, which has been shown in the previous section to affect the therapy significantly. The post-stroke damage to the blood-brain barrier can vary depending on whether the treatment is successful. It is also determined by the timing of clinical treatment and the collateral score of the patients [45].

To investigate the optimal treatment of patients with different BBB damage levels, values is varied while the rest of parameters are uniformly random. 2000 virtual patients are generated for each. Similarly, to exclude patients with initial ICP lower than 20 mmHg, only the cases with initial ICP values over 25 mmHg are included (with a fraction of around 5%, 25%, 50%, 62%, respectively).

In Fig. 7a, the efficiency of treatment rises with the, from a mean of around 71.2 9.8% to 93.3 6.8%. Meanwhile, the distributions of time ratio in the patient groups show similar patterns. To investigate the proper dosage for the patient groups, bar charts for the patient number who had ineffective osmotherapy are shown. Fig. 7b&c presents the number of patients who had time ratio of 0 and time ratio below 25%, respectively. Here, zero time ratio means that 20mmHg is not achieved, and below 25% time ratio means 20mmHg is achieved less than an hour. Overall, an osmotic pressure of 2000 Pa helps reach a ICP below 20 mmHg and 2500 Pa ensures that most patients are treated effectively.

**Figure 7:**
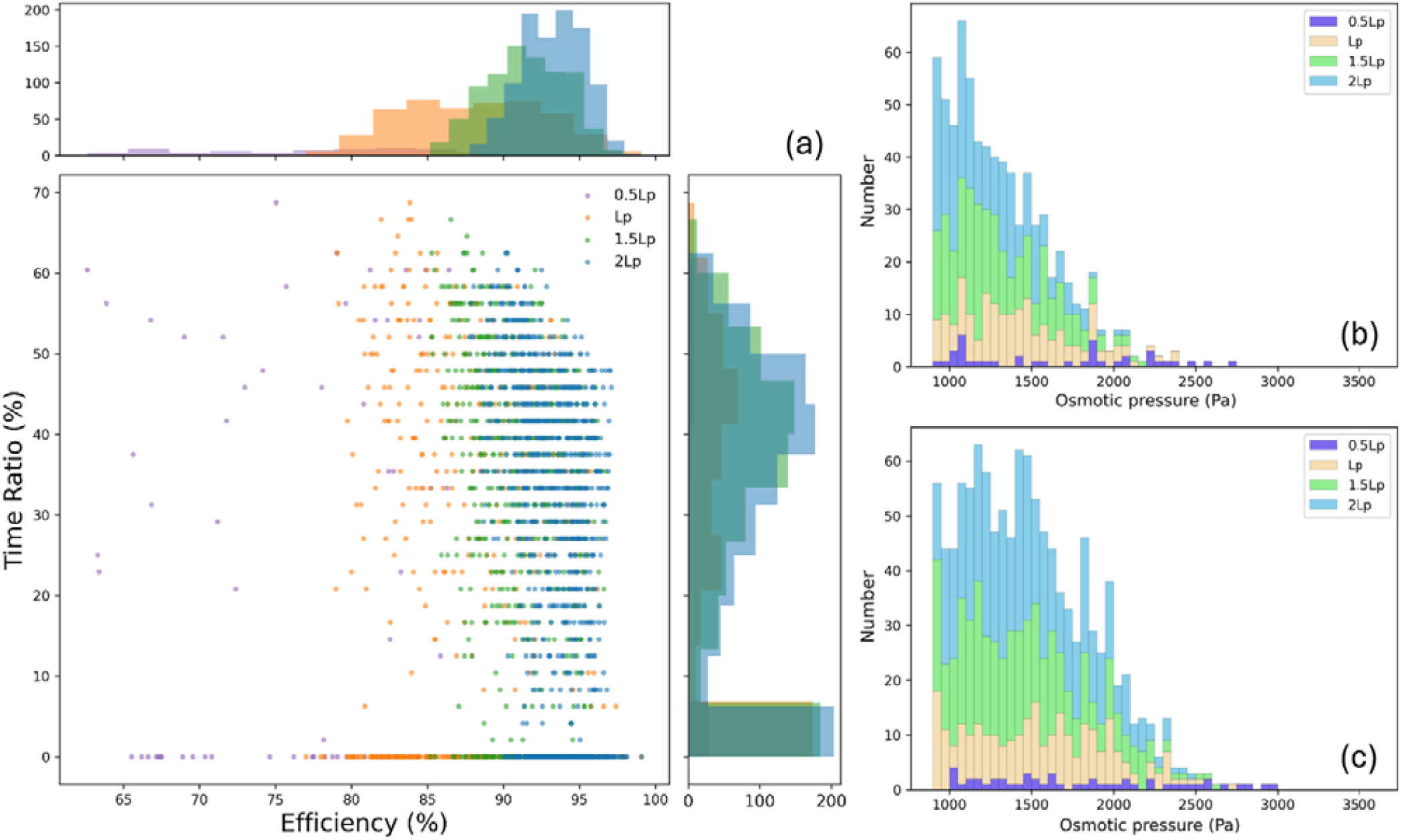
(a) The efficiency and time ratio under different values (0.5, 1, 1.5, 2 times the baseline) and doses of osmotic agent at each injection. (b) Patients are categorised by the maximum osmotic pressure to show patient number with zero time ratio. (c) Patients are categorised by the maximum osmotic pressure to show patient number with below 25% time ratio.

### E. The Optimization of Drug Doses for Various Age Groups

Here, we attempt to relate the model parameters with certain patient groups. In clinical settings, the age of patients is an important factor to consider as ageing changes not just the biomedical conditions of the patient but also the mechanical properties of the human brain. According to previous studies, ageing is related to a softening of brain tissue [31, 46], increased blood vessel leakage [43, 47], a decrease in water content in the brain [48-50], a decrease in tissue conductivity [48]. These indicate a lower tissue hydraulic conductivity, a higher vessel leakage rate, larger storage factor in the elder patients. Meanwhile, the mean arterial pressure (MAP) is not found to be statistically different between young and elderly patients [51]. The previous studies on these parameters are summarised in Table B1.

Although there is a decrease in water content in the brain tissue and a reduced hydraulic conductivity in rats, quantitative studies on human brain tissue hydraulic conductivity are rare. In [52, 53], a decrease in the CSF production rate (around half CSF production rate in the 70s compared to the 30s), whilst CSF production rate is found not age-dependent but rather individual-dependent in a more recent study [54]. It is therefore inferred from the data [52-54] that the 50s has 1.5 the hydraulic conductivity compared to the 90s as a result of the CSF production rate decrease in the elderly’s brains. Meanwhile, the tension test of the brain tissue shows a doubled tension shear modulus in the 50s subjects compared to the 90s subjects [31] and there has been shown a water content decrease with ageing [50]. This indicates a larger storage factor in the 50s. Additionally, 1.5 times the blood vessel wall permeability is implemented to reflect the 1.5 times difference in vessel wall leakage between the 50s and 90s populations [43], whereas the arteriole blood pressure at the pial surface is considered an approximation to mean arterial pressure (MAP).

To investigate the effects of ageing, patients between 50 to 90 years old are studied and 2000 virtual patients are generated for each group. The vessel wall permeability is given a random number between 0.5-1.33 the baseline value for the 50 years group, and these randomly generated values are multiplied 1.5 to represent the 90 years group. The storage factor is 0.5-1 baseline value in the 50s and doubled in the 90s. Tissue hydraulic conductivity is given a random number between 0.5-1.33 the baseline value in the 90s and 0.75-2 in the 50s age group. In the meantime, the arteriole blood pressure values are generated randomly between 80 to 100 mmHg for both age groups. The efficiency and time ratio for the evaluation of osmotherapy episodes are shown in Fig. 8.

**Figure 8:**
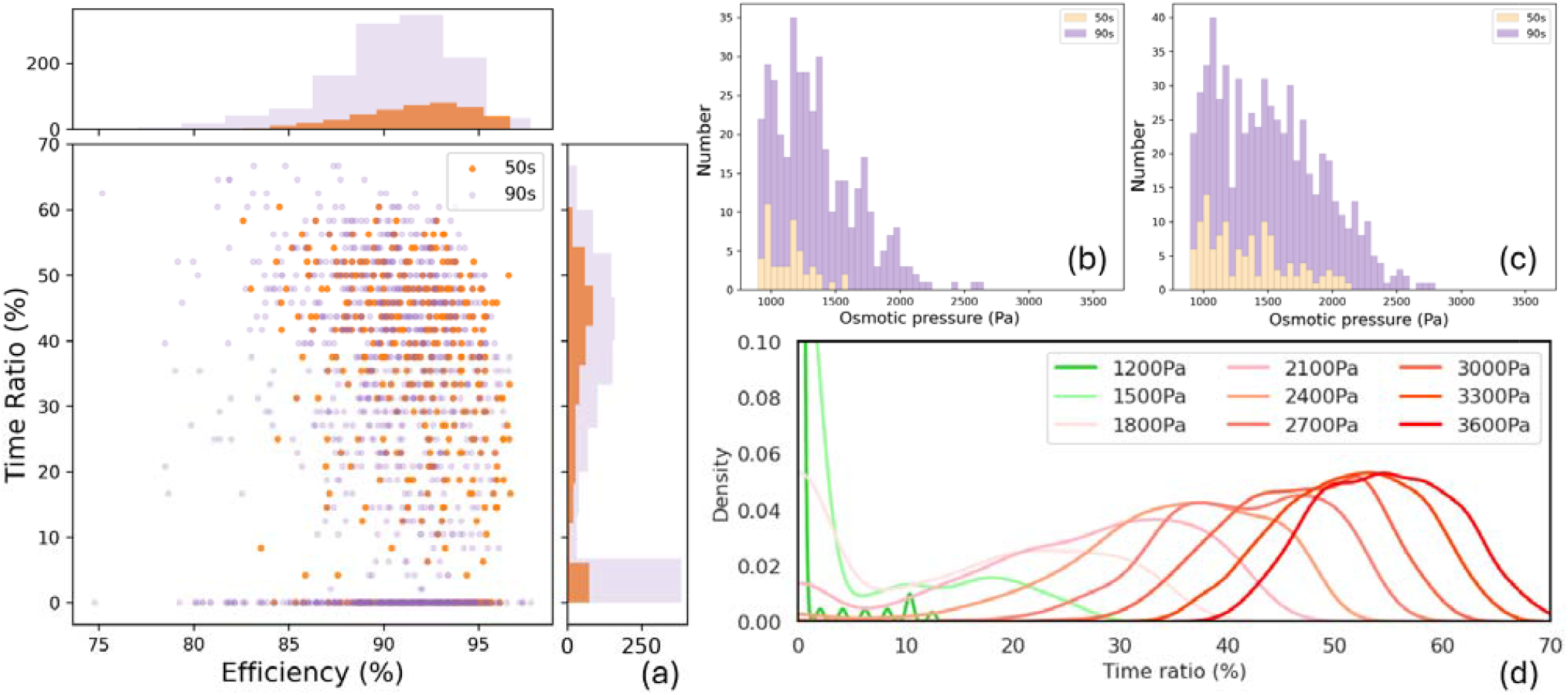
(a) The efficiency and time ratio of the 50s and the 90s age groups. (b) Patients are categorised by the maximum osmotic pressure to show patient number with zero time ratio. (c) Patients are categorised by the maximum osmotic pressure to show patient number with below 25% time ratio. (d) histogram of the time ratio for the 90s under various maximum osmotic pressure.

In the results, it can be seen that the 90s has a more than three times incidence of developing oedema beyond 25 mmHg (Fig. 8a). The efficiency and time ratio of the 50s and 90s age group are similar, with a higher risk of ineffective (20 mmHg not achieved) in the 90s. meanwhile, Fig. 8b&c show that 1500-2000 Pa osmotic pressure is needed to effectively treat the 50 s, whereas 2000-2500 Pa is needed for the 90s. Finally, Fig. 8d shows the probability of the time ratio of the 90s treated with different doses. It is found that a dosage that creates osmotic pressure of 1500 Pa and below can be ineffective in most cases, but the effectiveness can rise significantly when a threshold is reached (around 2100 Pa).

## IV. Discussion

This study presents the first osmotherapy in-silico trial assisted by a DNN. In clinical settings, the treatment of brain oedema is usually based on the medical history and other characteristics of the patients, for example, the ages, sex, etc. These patient data have been widely used to predict the outcome and the severity of oedema. However, how the medical history and patient characteristics are correlated with the ICP and with other flow parameters remains unclear. In this study, a DNN is utilised to generate virtual patients and the flow parameters are related to patient groups of various levels of BBB damage and ages. To the best of our knowledge, this is the first attempt to correlate the clinical statistics to a FEM mathematical model. This helps tackle the challenge in osmotherapy studies as the ICP data can be rare in clinical settings. More importantly, this study bridges the gap between bioinformatics and biophysical laws by providing an osmotherapy model that is supported by both well-founded mathematical /mechanical theories and patient data.

The model generates simulation results that are in good match with the clinical data when compared with 8 oedema patients (average ages of 59.8 8.8) who underwent 22 episodes using 10% hypertonic saline (Fig. 2e). In section 3.2, the simulation results show that the initial ICP ranges from around 16 to 29 mmHg, which agrees well with the clinical ICP rise to [15, 30] mmHg [4]. Meanwhile, the effect of hypertonic saline on ICP lasts more than 2 hours in the model, the mean of maximum ICP drop 9.72 mmHg, and nadir values are reached between 15-115 mins after the therapy. This matches excellently with clinical studies, where the mean maximum ICP reduction is 10.1 [55], the nadir values are achieved at 15-120 min [55, 56], and the ICP usually remain decreased 1-5+ hours after infusion [8, 56]. Furthermore, the cases with 3600 Pa maximum osmotic pressure (20% hypertonic saline) generates a pressure drop of around 54.3% from initial ICP (Fig. 5), which agrees well with the retrospective studies [44, 57], where a 55.6% and around 50% pressure drop was found in 23.4% hypertonic saline treatment. Generally, the ICP curves generated from the study can match the clinical data in terms of the nadir, initial ICP values and the time these critical values are reached. It therefore guarantees that the in-silico trials are comparable to clinical data.

However, previous clinical studies majorly focus on the doses and types of osmotic agents and there is no existing study on patient stratification. This is because ICP measurement involves high risks and the patient samples are usually small. This makes it impossible to further stratify the patients in clinical studies. Here, in-silico trials can be a nice complementary tool to investigate osmotherapy in different patient groups. In the DNN predicted results, the initial ICP mainly depends on the and the hydraulic conductivity. In contrast, the effect of arteriole blood pressure is not as significant as a variation of ABP from 80 to 100 mmHg only leads to an around 2 mmHg difference in the initial ICP in the virtual patients. This indicates that fluid accumulation is highly dependent on the restriction of the blood-brain barrier on the fluid flow through the vessel wall. Meanwhile, the lowest ICP is strongly affected by the choice of storage factor (Fig. 5). As brain tissue with a small storage factor can respond to the osmotic pressure rise in the blood vessel in a shorter period of time, brains with a small storage factor also experience a faster pressure rebound after the osmotic pressure drops as the drug concentration starts dropping. Therefore, the storage factor of the brain tissue has multiple effects on the ICP curve. Interestingly, the virtual patient data show that a smaller storage factor can lead to a lower minimum ICP, larger time ratio and higher efficiency during an episode. This indicates that a small storage factor (saturated brain) is favourable to osmotherapy.

Furthermore, the AI-assisted in-silico trial is evaluated based on two criteria, the time ratio, and the efficiency. The model is used to study different patient groups, including patients with various levels of BBB damage and patients at different ages, namely the 50s and the 90s age group populations. In the patient groups with different l__p_ values, the efficiency varies whilst the time ratio shows a similar pattern among these groups. Differing from our previous study on osmotherapy [26], where the dose of the medication shows a strong influence on specific cases, only patient groups are considered in Fig. 7b&c. It is challenging to obtain the exact value of the parameters and it is therefore reasonable to consider an optimization on a population-averaged level. In Fig. 7b&c, the virtual patients show a 2000-2500 Pa osmotic pressure is sufficient to achieve ICP below 20 mmHg in most cases. Considering the side effects and the marginal ICP drop, higher doses are thus not proposed unless radical treatment is needed in clinical settings.

Finally, the simulation results present a higher possibility of osmotherapy failure in the 90s population and the virtual patient groups show that a larger dose is needed for the elderly (90s) population to guarantee a ICP below 20 mmHg (Fig. 8). Furthermore, the histogram of time ratio in the 90s show a threshold in the effectiveness in osmotherapy (Fig. 8d). Therefore, a failure in osmotherapy can have multiple reasons: ineffectiveness when using small doses below the threshold; and insensitivity of patients to certain types of osmotic agents [2, 30]. Therefore, the dosage needs to be sufficiently improved before another osmotic agent or other therapeutic strategies are considered.

### Limitations

It should be noted that there are several limitations to this study. Firstly, many parameters have been used in the multi-compartment models and it is challenging to determine some of the parameters from the existing experimental data. Due to the limited experimental data, there lacks comparison between the parameter values in different studies. As far as we know, [32, 33] are the only study on the tissue hydraulic conductivity conducted by one group. The tissue hydraulic conductivity value is thus chosen from the range of experimental data in [32, 33]. However, such value can depend on the experimental methods and brain tissue samples used, and therefore can be further examined when more experimental data become available in parallel studies. Secondly, the parameters including *K*_*W*_, *C*_*w*_ and *L*_*p*_ data for different age groups are not yet available in the oedema human brain and data from healthy brain were thus used as necessary [31, 43, 52-54]. Finally, the model only takes into account the difference in the brain mechanical properties among patient groups. Other factors such as biochemistry and pathology are not considered here.

## V. Conclusions

In conclusion, this is the first study combining the conventional FEM model with AI methods to investigate osmotherapy on a quasi-population level. The model bridges the gap between biophysical laws and bioinformatics to provide a novel approach to studies various patient groups in osmotherapy. Once more accurate mechanical properties and physiological parameters of the brain become available. The model will be able to provide a more accurate prediction of clinical outcomes and thus improve the clinical treatment of brain oedema.

## Appendix A Derivation of Fluid Filtration through Capillary Wall

Here, we employ the filtration theory and the Donnan effect, which has been used to model brain tissue in our previous studies [26], to derive *S*_*cw*_, i.e., fluid filtration through the damaged BBB. The filtration flux, J_*VA*_, can be described by the following filtration equation as shown below.

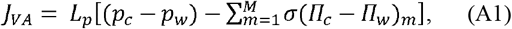

where *L*_*p*_ is the hydraulic permeability of the capillary wall, m is the different solutes, *p*_*i*_ is the hydrostatic pressure with subscripts that represent capillary blood pressure and interstitial fluid pressure. Meanwhile, Π_*c*_ is the osmotic pressure for each water-soluble solute present in the blood plasma and interstitial fluid. σ is the reflection coefficient that varies from 0 to 1, where a value of 1 means that no solute can flow through the BBB. Meanwhile, the difference in osmotic pressure can be described by the equation:

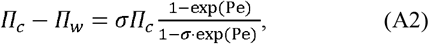

where Pe is the Péclet number. Here the Péclet number is much smaller than one [26] and for a fixed composition of the filtration model, we thus have

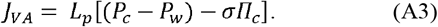

Meanwhile, the fluid transfer between capillary network and interstitial space *S*_*cw*_ can be given as

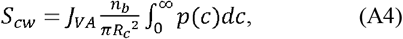

where *n*_*b*_ is the volume fraction of blood vessel in a unit volume of brain tissue, *R*_*c*_ is vessel radius and p(*c*), at each point in space over the distribution of vessel circumferences. By assuming that the perimeter of blood vessels remains constant, we have

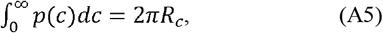

Substituting Equations A3 and A5 into Equation A4, *S*_*cw*_ can therefore be written as:

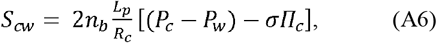

Note that, for osmotherapy, we introduce an additional term to include the effects of saline:

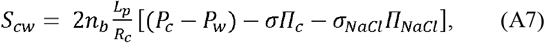

*L*_*p*_ in the normal BBB can increase by more than 100 times in oedema. Due to the large difference between the BBB permeability in the damaged region and the healthy region, the flow from capillary blood to the interstitial space and the flow of interstitial fluid in healthy tissue is thus neglected [26].

## Appendix B Parameter Value Ranges

The ranges of the parameter values from the literature are listed in Table B1:

**TABLE B1.** Sources of parameter values in the 50s and 90s.

| Physiological Parameters |  |  |
| --- | --- | --- |
| Arteriole blood pressure [51] | 95±11 mmHg in the 50s | 98±11 mmHg in the 90s |
| Tension shear modulus [31] | Around 500 Pa in the 50s | Around 250 Pa in the 90s |
| CSF production rate [52] | 0.41±0.24 in the 30s | 0.19±0.07 in the 80s |
| Vessel wall leakage rate [43] | 7.5 min<sup>-1</sup> in the 50s | 12 min<sup>-1</sup> in the 90s |

An around two times difference are shown in l__p_ and tension shear modulus between the 50s and 90s groups. Although the vessel leakage rate is measure in healthy brains, the value is simply taken to be around 1.5 times in oedema patients as there is not yet any relevant dataset.

## Appendix C Model Parameters

**TABLE C1.** Model Parameters.

| Parameter Values |  |  |  |
| --- | --- | --- | --- |
| $c_b$ | $1.59 \times 10^{-3} \text{ Pa}^{-1}$ | $R_c$ | $5 \times 10^{-6} \text{ m}$ |
| $c_w$ | $3.08 \times 10^{-4} \text{ Pa}^{-1}$ | $\sigma$ | 0.35 |
| $K_a$ | $1.234 \text{ mm}^3 \text{ s} \cdot \text{kg}^{-1}$ | $\sigma_{NaCl}$ | 0.062 |
| $K_c$ | $4.28 \times 10^{-3} \text{ mm}^3 \text{ s} \cdot \text{kg}^{-1}$ | $\Pi_c$ | 2445 Pa |
| $K_v$ | $2.468 \text{ mm}^3 \text{ s} \cdot \text{kg}^{-1}$ | $\omega_{ac}^g$ | $1.326 \times 10^{-6} \text{ Pa}^{-1} \text{ s}^{-1}$ |
| $K_w$ | $3.6 \times 10^{-3} \text{ mm}^3 \text{ s} \cdot \text{kg}^{-1}$ | $\omega_{ac}^w$ | $5.22 \times 10^{-7} \text{ Pa}^{-1} \text{ s}^{-1}$ |
| $L_p$ | $3.0 \times 10^{-11} \text{ m/s} \cdot \text{Pa}$ | $\omega_{cv}^w$ | $4.641 \times 10^{-6} \text{ Pa}^{-1} \text{ s}^{-1}$ |
| $n_b$ | 0.03 | $\omega_{cv}^g$ | $1.828 \times 10^{-6} \text{ Pa}^{-1} \text{ s}^{-1}$ |

